# Temporal windows of perceptual organization: Evidence from crowding and uncrowding

**DOI:** 10.1101/2025.07.14.664690

**Authors:** Alessia Santoni, Luca Ronconi, Jason Samaha

## Abstract

Organizing visual input into coherent percepts requires dynamic grouping and segmentation mechanisms that operate across both spatial and temporal domains. While crowding disrupts target perception when nearby elements fall within the same spatial pooling window, specific flanker configurations can alleviate this effect through Gestalt-based grouping, a phenomenon known as uncrowding. Here, we examined the temporal dynamics underlying these spatial organization processes using a Vernier discrimination task. In Experiment 1, we varied stimulus duration and found that uncrowding emerged only after 160 ms, suggesting a time-consuming process. In Experiment 2, we manipulated the stimulus onset asynchrony (SOA) between the target and flankers. We found that presenting good-Gestalt flankers briefly before the target (as little as 32 ms) significantly boosted uncrowding, even in the absence of temporal overlap between the two stimuli. This effect was specific to conditions in which flankers preceded the target, ruling out pure temporal integration and masking accounts. These findings suggest that spatial segmentation can be dynamically facilitated when the temporal order of presentation allows grouping mechanisms to engage prior to target processing. Moreover, the observed time course indicates that segmentation is not purely feedforward, particularly for stimuli that are likely to recruit higher-level visual areas, pointing instead to the involvement of recurrent or feedback processes.

## INTRODUCTION

In order to organize incoming sensory input, the visual system needs to identify and group elements across space, as well as integrate information across time. This dual challenge is central to defining coherent and meaningful perceptual units, yet the mechanisms that support it remain only partially understood.

In the spatial domain, the presence of nearby elements in the visual scene can impair target perception, a phenomenon known as crowding. Crowding is a pervasive feature of visual processing, occurring across different levels of stimulus complexity, from simple Gabor patches to alphanumeric characters and natural scenes (e.g., Bernard & Chung, 2011; Ringer et al., 2021; Ronconi et al., 2016; Tanriverdi & Cornelissen, 2024). Previous research has identified object spacing as the key factor driving crowding: when the distance between target and flankers falls below an observer’s critical spacing at that retinal location, crowding is likely to occur. This principle, originally described by Herman Bouma in 1970, has since become known as “Bouma’s law” (Bouma, 1970; Pelli & Tillman, 2008). Additionally, the strength of crowding is shaped by flanker-target similarity, with target identification errors often reflecting either a bias toward flanker features or mislocalization of the target relative to the flankers (Bernard & Chung, 2011; Põder & Wagemans, 2007; Zahabi & Arguin, 2014). This and other evidence have contributed to the view that crowding reflects a perceptual bottleneck, arising from feature integration or pooling within spatial regions defined by receptive field sizes in the feedforward visual hierarchy (for a comprehensive characterization of models of crowding refer to Pelli, 2008).

This account of crowding has been challenged in the past decade by a notable exception in which specific flanker configurations alleviate the otherwise detrimental effects of crowding. This effect, known as *uncrowding*, is often attributed to Gestalt grouping principles, whereby flanking elements are perceptually organized into unitary structures based on their features (Malania et al., 2007; Manassi et al., 2012, 2013; Sayim et al., 2010). The spatial arrangement of flankers determines whether they are perceived as part of a unified object with the target or as separate elements, affecting the degree of crowding accordingly (Livne & Sagi, 2007; Saarela et al., 2010; Tiurina et al., 2022). Unlike crowding, which has been often considered an early, feedforward limitation, uncrowding may reflect a time-dependent process of perceptual organization emerging through recurrent or integrative mechanisms over time (Herzog et al., 2015; Manassi & Whitney, 2018).

In the temporal domain, crowding effects can still be observed when flankers are not presented simultaneously to the target. For example, Huckauf and Heller (2004) manipulated the stimulus onset asynchrony (SOA) between target and flanker letters and found that strong crowding effects emerged at short temporal delays, and persisted even when target and flankers were separated by longer time-windows of ±150 ms. However, a precise temporal characterization of uncrowding is still lacking. Sayim and colleagues (2014) investigated the interaction between backward masking (i.e., when the target precedes the flankers) and (un)crowding by manipulating both the appearance of the flankers and the SOA between target and flankers. They found that flankers can produce type-B backward masking, with the greatest performance impairments occurring at SOAs of approximately 40-60 ms, depending on stimulus characteristics. Notably, the study did not reveal any perceptual advantage when flankers formed a good Gestalt (e.g., cuboids) compared to typical crowding-inducing flankers (e.g., vertical lines), suggesting that Gestalt-based representations may require additional time to emerge and be the results of higher-level feedback mechanisms (Sayim et al., 2014). This interpretation is also supported by recent results by Morea and colleagues (2025), who showed that while uncrowding effects typically emerged only after 160 ms under standard simultaneous presentation, even a brief 20 ms preview of the good-Gestalt flankers prior to the full flanker-target display significantly improved Vernier discrimination performance.

The aim of the present study is to investigate temporal dynamics underlying spatio-temporal organization in visual perception. To characterize the temporal dynamics of crowding and uncrowding, we conducted two experiments using a Vernier discrimination task. In the first study, we manipulated stimulus duration while presenting flankers known to elicit either crowding or uncrowding effects. In the second study, we introduced a temporal delay between the flankers and the target Vernier stimulus, such that flankers (inducing either crowding or uncrowding) could appear before or after the target. This design allowed us to systematically map both the directionality and timing of uncrowding effects. Our results first replicated previous findings showing that grouping effects typically emerge relatively slowly. However, this delay was drastically reduced when flankers were presented before the target. We interpret these findings in light of feedback or recurrent mechanisms, suggesting that early presentation of the flankers pre-activates these processes, leading to a marked reduction in the time required for the uncrowding benefit to emerge.

## METHODS

### Participants

24 participants were recruited from the University of California, Santa Cruz community. Two participants were excluded due to uncorrected visual impairments, resulting in a final sample of 22 participants (mean age = 22 years, range: 18-35 years). The final sample included 16 individuals who identified as female, 5 as male, and 1 as non-binary. Participants received university research credits as compensation, if interested. The study was approved by the Institutional Review Board at the University of California, Santa Cruz. All participants performed two experiments, which order was counterbalanced across participants.

### Apparatus and stimuli

Participants were seated in a dimly lit room at a viewing distance of 75 cm, maintained with a chinrest, and completed two tasks, each lasting approximately 25 minutes. Stimuli were presented through PsychToolbox 3 (Brainard, 1997) for MATLAB (The MathWorks Inc., 2022) on a black background on a VIEWPixx/EEG monitor with a 120 Hz refresh rate. Stimulus luminance was set to 80 cd/m^2^. In a Vernier discrimination task, Vernier targets consisted of two vertical lines, with a slight horizontal offset in the lower line. Following an intertrial interval (ITI) of 500 ms (±250 ms jitter), Vernier stimuli were randomly presented either to the left or right of the central fixation dot. The center of the stimulus was positioned 4 degrees of visual angle (DVA) from fixation, and participants were instructed to maintain central fixation throughout the trial. The two vertical lines composing the Vernier stimulus measured 40 arcminutes (′) in length each and were separated by a vertical gap of 4′. The horizontal offset between the two lines was set to 1′, and offset direction was randomized across trials. Across three experimental blocks, Vernier stimuli were presented either in isolation (baseline condition) or flanked by additional elements (illustrated in Figure 1A). Flankers, when present, were presented symmetrically to the Vernier stimulus along the horizontal axis at a distance of 16′. Flankers could be either vertical lines (44′ in length) or rectangular flankers (44′ tall × 116′ wide), previously shown to elicit crowding and uncrowding effects, respectively (Manassi et al., 2012). Participants were instructed to indicate the direction of the offset of the lower line relative to the upper line by pressing “M” (right) or “Z” (left) on the keyboard.

**Figure 1.**
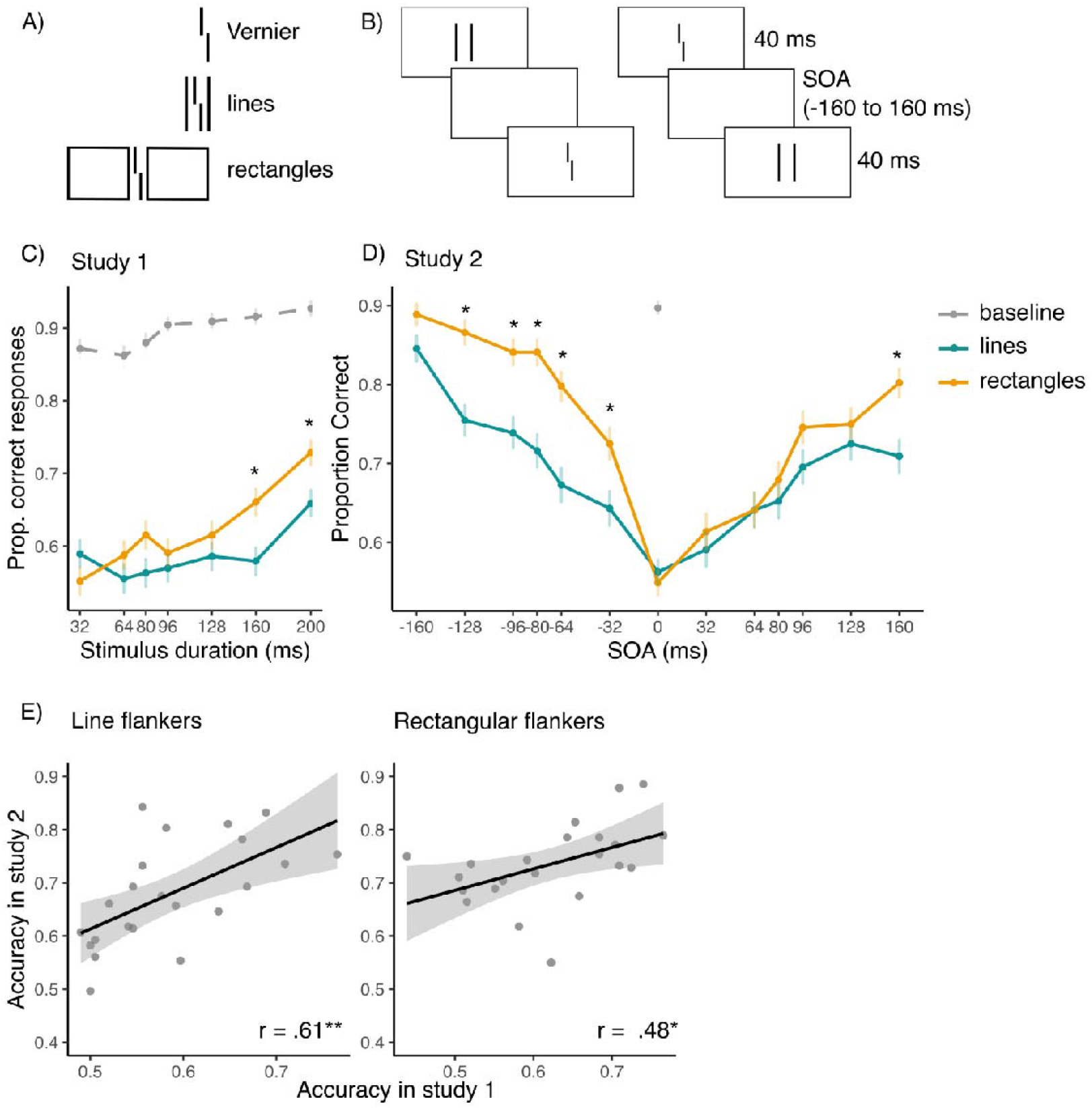
A) Stimuli used in the Vernier discrimination task; Vernier stimuli were either presented in isolation (baseline condition) or flanked by lines or rectangles across separate blocks of trials. B) Schematic representation of Experiment 2. A temporal delay was introduced between the flankers and the Vernier target, such that flankers either preceded (left panel) or followed (right panel) the Vernier. Stimuli are enlarged for illustrative purposes. On each trial, stimuli were randomly presented to the left or right of a central fixation point. C-D) Results from experiment 1 and 2 show that the uncrowding effect emerges around 160 ms when stimuli are presented simultaneously, but at a SOA of 32 ms when flankers precede Verniers. Asterisks represent *p*-values from t-tests comparing accuracy for flanker shape (lines vs. rectangles). Error bars represent standard error of the mean (SEM). E) Pearson’s correlations between accuracy in study 1 and 2, across conditions (left panel: line flanker blocks, right panel: rectangular flanker blocks). Dots represent individual subjects; grey shaded areas represent 95% confidence intervals. * *p*_*FDR*_ < .05, ** *p*_*FDR*_ < .01.

### Procedure

#### Experiment 1

Participants were asked to report the offset direction of the Vernier stimulus, while stimulus duration was manipulated along the following values: 32, 64, 80, 96, 128, 160, or 200 ms. Vernier and flanker stimuli were always presented simultaneously. The task included a total of 588 trials: 196 baseline trials and 196 trials for each flanker shape (lines and rectangles).

#### Experiment 2

Participants performed a Vernier discrimination task where the duration of both the Vernier and flanker stimuli was fixed at 40 ms. However, target and flanker stimuli were either presented simultaneously (i.e., a SOA of 0 ms) or with a variable SOA, randomly selected from ±32, ±64, ±80, ±96, ±128, or ±160 ms (Figure 1B). Negative SOA values indicate trials in which the flankers preceded the Vernier stimulus, whereas positive SOA values indicate trials in which the flankers followed the Vernier stimulus. The task included a total of 616 trials: 56 baseline trials and 280 trials for each flaker shape (lines and rectangles).

For both experiments, participants completed three separate blocks of trials: one for the baseline condition (no flankers), one with line flankers, and one with rectangle flankers. Block order was counterbalanced across participants. Before each experiment, participants completed 20 practice trials with feedback to familiarize themselves with the procedure.

### Statistical analyses

For experiment 1, we investigated the effect of stimulus duration on Vernier discrimination. With this aim, stimulus duration (seven levels: 32, 64, 80, 96, 128, 160, and 200 ms) and flanker shape (lines vs. rectangles) were input as within-subject factors in a two-way repeated-measures ANOVA on accuracy data. For experiment 2, a two-way repeated-measures ANOVA was conducted on accuracy data to examine the interaction between flanker shape (lines vs. rectangles) and stimulus SOA (twelve levels: ±32, ±64, ±80, ±96, ±128, ±160 ms). Sphericity was assessed using Mauchly’s test, and Greenhouse-Geisser correction was applied where necessary. For each ANOVA, post hoc t-tests were performed to further investigate the relationship between flanker shape and either stimulus duration or SOA. In the second experiment, additional post hoc t-tests were conducted to compare accuracy at same-magnitude positive and negative SOAs, for each flanker shape separately. False Discovery Rate (FDR; Benjamini & Hochberg, 1995) was used to control for the family-wise type-I error rate across multiple comparisons. Accuracy data from the baseline condition were not included in the ANOVAs; instead, they were examined to ensure that participants performed above chance level. Finally, Pearson’s correlation analyses were conducted to assess whether individual accuracy scores (averaged for stimulus duration/SOA level) based on flanker shape were related across the two tasks.

## RESULTS

### Experiment 1

Results from experiment 1 are depicted in Figure 1C. A two-way repeated-measures ANOVA was conducted to examine the effects of stimulus duration (seven levels: 32, 64, 80, 96, 128, 160, and 200 ms) and flanker shape (lines vs. rectangles) on Vernier discrimination accuracy. The analysis revealed a significant main effect of stimulus duration, *F*(3.47, 72.80) = 8.12, *p* < .001, and a significant main effect of flanker shape, *F*(1, 21) = 6.63, *p* = .018. Importantly, the analysis showed a significant interaction between stimulus duration and flanker shape, *F*(4.60, 96.60) = 2.51, *p* = .039, suggesting that the effect of duration on accuracy differed between the two flanker shapes. Post hoc pairwise comparisons conducted between the two flanker shapes at each stimulus duration revealed significantly higher accuracy in the presence of rectangular flankers compared to flanker lines at 160 ms, *t*(21) = -2.77, *p*_*FDR*_ = .040, and 200 ms, *t*(21) = -2.83, *p*_*FDR*_ = .040. No significant differences were found at shorter durations (32–128 ms; all *ps* > .08).

### Experiment 2

Results from experiment 2 are depicted in Figure 1D. The repeated-measures ANOVA on flanker shape (lines vs. rectangles) and SOA (twelve levels: ±32, ±64, ±80, ±96, ±128, ±160 ms) revealed a significant main effect of SOA, *F*(6.87, 144.24) = 32.96, *p* < .001, and a significant main effect of flanker shape, *F*(1, 21) = 22.88, *p* < .001. A significant interaction between SOA and flanker shape was found, *F*(7.69, 161.64) = 3.00, *p* = .004. Post hoc tests comparing accuracy for flanker shapes at each SOA revealed significantly higher accuracy for rectangular flankers at negative SOAs of -128 ms (*t*(21) = -3.99, *p*_*FDR*_ = .003), - 96 ms (*t*(21) = -3.70, *p*_*FDR*_ = .004), -80 ms (*t*(21) = -4.65, *p*_*FDR*_ < .001), -64 ms (*t*(21) = -3.38, *p*_*FDR*_ = .006), and -32 ms (*t*(21) = -3.37, *p*_*FDR*_ = .006), as well as at the positive SOA of 160 ms (*t*(21) = -4.64, *p*_*FDR*_ < .001). No significant differences were found at 0 ms or positive SOAs up to 128 ms (*p*_*FDR*_ > .10). Post hoc t-test revealed that Vernier accuracy significantly differed between negative and positive SOAs of the same magnitude across all SOA levels when rectangular flankers were presented, at 160 ms (*t*(21) = 3.82, *p*_*FDR*_ = .002), 128 ms (*t*(21) = 3.57, *p*_*FDR*_ = .002), 96 ms (*t*(21) = 3.57, *p*_*FDR*_ = .002), 80 ms (*t*(21) = 6.81, *p*_*FDR*_ < .001), 64 ms (*t*(21) = 4.50, *p*_*FDR*_ = 0.001), and 32 ms (*t*(21) = 3.21, *p*_*FDR*_ = 0.004). The same post hoc t-test performed when line flankers were presented only revealed a significant difference at 160 ms (*t*(21) = 5.68, *p*_*FDR*_ < .001, all other *p*s > .15).

### Correlations

Accuracy scores averaged for stimulus duration/SOA level across the two experiments were positively correlated for both line flanker blocks (*r*(20) = 0.61, *p* = .003) and rectangular flanker blocks (*r*(20) = 0.48, *p* = .025), indicating that the two tasks successfully measured related processes (Figure 1F).

## DISCUSSION

Perception depends on dynamic grouping and segmentation processes that unfold across both space and time. Here, we extended previous work by examining the time course of spatial segmentation using a Vernier discrimination paradigm known to produce strong crowding and uncrowding effects depending on the flanker configuration. Rectangular flankers, in particular, have been shown to alleviate crowding compared to simple vertical lines, as they are perceptually grouped into a single unit under Gestalt principles (Herzog et al., 2015; Manassi et al., 2012). While earlier studies have largely focused on how flanker properties influence spatial grouping (see, e.g., Manassi et al., 2012, 2013), our focus was on when these effects emerge during stimulus processing. In our first experiment, we mapped the timeframe of the uncrowding benefit by varying stimulus duration across a broad range (20 to 200 ms) and found that uncrowding emerged only when the stimulus was presented for at least 160 ms. This aligns with recent findings by Morea et al. (2025), who reported uncrowding at 160 ms but not at 20 ms, though their study included only two durations. By sampling more densely across time, our results confirm and extend their observation, suggesting that spatial segmentation and grouping build up over time, likely via either recurrent or feedback processing (Herzog et al., 2015). However, a difference from previous studies is that in our paradigm, stimulus location was randomized across hemifields, minimizing stimulus expectancy and preparatory spatial attention. Despite this, we observed that the same temporal window was necessary for uncrowding to emerge, indicating that the time course of spatial segmentation is likely not contingent on stimulus location predictability.

While our first experiment confirmed that spatial organization unfolds over relatively long timescales, our second experiment demonstrated that when flankers precede the target, the uncrowding effect emerges more rapidly. In this experiment, we systematically varied the SOA between the Vernier and the flankers, both presented for 40 ms, across a broad temporal range. First, we showed that even when flankers and Verniers are not presented simultaneously, crowding and uncrowding effects are still observed, demonstrating the temporal extent of spatial segmentation mechanisms. Furthermore, we found uncrowding benefits emerging at SOAs as early as 32 ms and persisting up to 128 ms when flankers preceded the target. Beyond this point, the temporal separation between the two stimuli was large enough that the temporal crowding effect was minimal to begin with. At an SOA of -32 ms, assuming temporal integration between flankers and Vernier, the total visual stimulation amounts to 72 ms, about half the presentation duration expected to elicit uncrowding under the condition of our first experiment. This suggests that Gestalt-based uncrowding can emerge much faster than previously thought when there is a brief temporal offset between the flanker and target.

Moreover, while an SOA of -32 ms involved an 8 ms temporal overlap between flankers and the Vernier, similar uncrowding effects were also observed at longer negative SOAs without any overlap (i.e., SOAs < -64 ms). For example, even at an SOA of -128 ms, corresponding to an 88 ms gap between flanker offset and Vernier onset, significant uncrowding still occurred. This highlights the temporal stability of Gestalt grouping, which can alleviate crowding of subsequent targets even after a substantial delay.

The task manipulation introduced in this second experiment allows us to investigate Temporal Integration Windows (TIWs) across different perceptual conditions; for example, we show that uncrowding stimuli define a narrower TIW as compared to crowding stimuli (see the width of the curve in Figure 1D). However, it is important to note that the observed uncrowding benefits cannot be solely attributed to temporal integration mechanisms. Indeed, this effect was specific to negative SOAs (i.e., when flankers preceded the target), implicating rapid preview mechanisms that extend beyond simple temporal integration of the two stimuli. If temporal integration were the only mechanism at play, the uncrowding benefit should have been comparable for positive and negative SOAs of equal magnitude.

On the other hand, our results are also only partially compatible with masking mechanisms. Asymmetries between forward and backward masking are well-documented, with backward masking typically exerting a stronger disruptive effect on perception (Enns & Di Lollo, 2000). However, our findings show that crowding strength was comparable for positive and negative SOAs of equal magnitude when presenting crowding stimuli (with the expectation of the SOA equal to 160 ms). According to the masking account, backward masking effects are often interpreted as arising from temporally delayed feedback from higher-to lower-level visual processes, which can disrupt the perception of the target stimulus (Di Lollo et al., 2000). In line with this idea, Huckauf and Heller (2004) showed that crowding was reduced when targets were followed by letter-like nonletters as opposed to letter flankers. They attributed this effect to reduced top-down activation: since letter-like nonletters engage the letter level more weakly, they generate less feedback to early visual areas, thereby weakening the crowding effect. Additional results investigating backward masking and stimulus configurations come from Sayim et al. (2014), where authors reported stronger masking at SOAs around 40-60 ms (i.e., type-B masking) regardless of flanker configuration (i.e., flankers could be either lines or cuboids). This lack of uncrowding effect for backward masking is coherent with our results at positive SOAs, further corroborating the idea that Gestalt representations might operate on feedback processes that require additional processing time (Wagemans et al., 2012). However, in our study, performance was maximally impaired at short and zero SOAs, rather than intermediate SOAs as in Sayim et al. (2004).

Taken together, we suggest that when flakers form good Gestalts, spatial grouping processes can rapidly segment flankers from targets and do so even in the absence of temporal overlap between the stimuli. It has been previously hypothesized that crowding occurs when stimuli fall within the same processing stage (Manassi & Whitney, 2018). In this view, pre-activating relevant Gestalt structures may engage top-down/lateral spatial segmentation mechanisms that facilitate subsequent target processing. In contrast, when the target preceded the flankers, no such advantage was observed, further arguing against a purely masking-based account.

Importantly, our conclusions about the involvement of feedback or recurrent mechanisms are likely to apply specifically to the relatively complex flanker configurations used in our paradigm, which presumably engage higher-level visual areas responsible for spatial grouping across objects and shapes. It remains an open question whether similar temporal dynamics would be observed with simpler stimuli that rely predominantly on early visual processing stages, where crowding might still arise from more purely feedforward mechanisms.

Finally, it is possible that exogenous attention contributed to the behavioral improvement observed at negative SOAs compared to their positive counterparts. Previous studies have shown that both endogenous and exogenous attention can modulate the strength of crowding (Gong et al., 2024; Kewan-Khalayly & Yashar, 2022; Yeshurun & Carrasco, 1999). However, the timeline of our fastest stimuli is not entirely compatible with mechanisms of exogenous attention, which tend to emerge around 100 ms after cue onset (Carrasco, 2011; Egeth & Yantis, 1997). At any rate, it is reasonable to assume that any potential attentional effects in our paradigm were equally present in the two flanker configurations and therefore cannot fully account for the uncrowding effects observed here.

## CONCLUSIONS

Our results demonstrate that spatial segmentation, though relatively slow under standard uncrowding conditions, can be significantly accelerated when uncrowding flankers are presented prior to the target stimulus. We interpret this finding as evidence that a rapid presentation of the flankers, even for just tens of milliseconds, facilitates the emergence of Gestalt-based benefits by initiating grouping mechanisms that have a sustained impact on subsequent target perception. In turn, these results offer further support for the idea that spatial segmentation and grouping can be mediated by recurrent or feedback processing in the case of relatively complex configurations that likely engage higher-level visual areas.

## ACKNOWLEDGMENTS

We would like to thank Michael Herzog and Martina Morea for the helpful discussions. We also thank Emily Lincoln, Vrishab Nukala and Gaia Minari for their help in data collection. The present work was performed by A.S. in fulfilment of the requirements for obtaining the PhD degree at Vita-Salute San Raffaele University, Milan, Italy.

